# Integrated genetic and methylomic analyses identify shared biology between autism and autistic traits

**DOI:** 10.1101/493601

**Authors:** Aicha Massrali, Helena Brunel, Eilis Hannon, Chloe Wong, iPSYCH-MINERvA Epigenetics Group, Simon Baron-Cohen, Varun Warrier

**Affiliations:** Autism Research Centre, University of Cambridge, UK; University of Exeter Medical School, University of Exeter, RILD Building, Level 4, Barrack Rd, Exeter, UK; King’s College London, Institute of Psychiatry, Psychology and Neuroscience, De Crespigny Park, London, UK; Cambridgeshire and Peterborough National Health Service Foundation Trust, Cambridge, UK

**Author notes:** These authors contributed equally to this manuscript (Aicha Massrali, Helena Brunel, Varun Warrier). Correspondence to: Aicha Massrali, Helena Brunel, or Varun Warrier.

## Abstract

Previous studies have identified differences in DNA methylation in autistic individuals compared to neurotypical individuals. Yet, it is unclear if this extends to autistic traits – subclinical manifestation of autism features in the general population. Here, we investigate the association between DNA methylation at birth (cord blood), and scores on the Social and Communication Disorders Checklist (SCDC), a measure of autistic traits, in 701 8-year olds, by conducting a methylome-wide association study (MWAS) using DNA methylation data from cord-blood. Whilst did not identify significant loci demonstrating differential methylation, we observe a degree of overlap between the SCDC MWAS and post-mortem brain methylation signature in autism. Validating this, we observe an enrichment for genes that are dysregulated in the post-mortem autism brain. Finally, integrating genome-wide data from more than 40,000 individuals and mQTL maps from cord-blood, we demonstrate that mQTLs of CpGs associated with SCDC scores at different P-value thresholds are significantly shifted towards lower P-values in a GWAS for autism. We validate this using a GWAS of SCDC, and demonstrate a lack of enrichment in a GWAS of Alzheimer’s disease. Our results highlight the shared cross-tissue epigenetic architecture of autism and autistic traits, and demonstrate that mQTLs associated with methylation changes in childhood autistic traits are enriched for common genetic variants associated with autism and autistic traits.

## Introduction

Autism is a neurodevelopmental condition characterized by social-communication difficulties, unusually restrictive, repetitive behaviour and narrow interests, and sensory difficulties^1,2^. The condition can be thought as a continuum, with autistic traits being normally distributed in the general population, and autism at the extreme end of the continuum^3–5^. Both autism and autistic traits are highly heritable^6–9^, with variation across the allelic spectrum associated with the condition^10–12^. Despite a significant SNP heritability, recent studies have demonstrated that the variance explained per SNP is small, suggesting a highly polygenic architecture^11,13^. None of the significant SNPs associated with autism result in predicted coding changes, suggesting that they regulate gene expression through other mechanisms^11,14^. For instance, a recent genome-wide association study (GWAS) of autism has identified an enrichment of GWAS signals in H3K4me1 histone marks, particularly in brain and neural cell lines^11,13^.

One potential mechanism through which common genetic variants can regulate gene expression is through DNA methylation. DNA methylation is partly heritable^15,16^, with approximately 40% of CpG sites having a significant genetic component^17^. Previous studies have investigated autism associated methylation signatures in both peripheral tissues^14,18,19^ and in the post-mortem brain^20–23^. While post-mortem brain is pertinent for a neurodevelopmental condition like autism, it is not readily accessible, and will be confounded by post-mortem effects on methylation patterns. Studies of methylation signatures in postmortem brains in autism have replicably identified differential methylation^20–23^. Further, they have demonstrated an enrichment for differentially methylated signatures in immune system, synaptic signalling and neuronal regulation^20,21,23^. In contrast, recent large-scale analysis of three different peripheral tissue datasets have not identified significantly differentially methylated CpG sites in cases compared to controls^14,19^. The lack of significant results in peripheral tissues may be attributable to small effect sizes, significant heterogeneity in both CpG methylation and autism.

Given the heritability of methylation, a few studies have integrated genetics and methylation marks to identify convergent signatures in autism. Andrews and colleagues demonstrated that autism associated GWAS loci are enriched for methylation QTLs (mQTLs) in foetal brain and blood, suggesting that at least some of the genetic loci associated with autism may contribute to the condition through differential methylation^24^. In line with this, Hannon and colleagues demonstrated that polygenic risk for autism is associated with differential methylation at birth^14^. While these studies have demonstrated a role for common genetic variants associated with autism and influencing methylation, to our knowledge no study has investigated if methylation of CpGs associated with autism or autistic traits are enriched for polygenic signatures of autism or autistic traits. One way to test this hypothesis is using mQTLs. We hypothesized that given methylation at CpGs is partly driven by genetics through mQTLs as evidenced by the significant heritability of methylation marks, then we would expect mQTLs of significant CpGs in a methylome-wide association study of autism or autistic traits to be enriched for lower P-values in a GWAS of autism or autistic traits.

To address these questions, we investigated the association of CpG methylation in cord blood using a measure of social autistic traits at age 8 (scores on the Social and Communications Disorder Checklist, or SCDC)^25^. The SCDC is phenotypically and genetically correlated with autism (r_g_ ~ 0.3)^5,25,26^, and polygenic scores from autism are associated with SCDC scores in the general population^26^. The advantage of using a continuous measure of autistic traits is that it captures the underlying variance better, and minimizes the heterogeneity due to different diagnostic criteria and practices used to diagnose autism. Further, the use of cord-blood CpGs minimizes (though, does not eliminate) reverse causation (where the phenotype influences DNA methylation), as the methylation of CpG sites is measured very early in life. To investigate how comparable an MWAS of an autistic trait is to other MWAS of autism and related phenotypes conducted across tissues and phenotypes, we investigated the overlap between the MWAS of SCDC and other MWAS of autism and communication-related traits in peripheral and post-mortem brain tissues. We further investigated if genes that are transcriptionally dysregulated in the post-mortem autism brain are enriched for methylation CpGs associated with SCDC. Finally, integrating GWAS data for autism from 46,350 individuals, we investigate if mQTLs of CpGs associated with SCDC scores at various P-value thresholds are significantly shifted towards lower P-values in the autism GWAS. We validate these results using a smaller GWAS for SCDC.

In summary, this study had two specific aims: 1. To investigate if an MWAS for autistic traits identifies significant CpG methylation and if it is comparable to MWAS of autism; 2. To investigate if genetic variants associated with autism and autistic traits are enriched in CpG sites associated with autistic traits at various P-value thresholds.

## Methods

### Study population

Participants were children from the Avon Longitudinal Study of Children and Parents (Children of the 90s)^27^, drawn from the Accessible Resource for Integrated Epigenetic Studies (ARIES, www.ariesepigenomics.org.uk)^28^. ALSPAC is a longitudinal cohort in which the participants were pregnant women in the Avon region in the UK. The initial cohort consists of 14,541 initial pregnancies and 13,988 children who were alive at the age of 1. In addition, children were enrolled in further phases. Details of the data available can be found on the online data dictionary here: http://www.bristol.ac.uk/alspac/researchers/access/. Written informed consent was obtained from the parent or the guardian and assent was obtained from the child where possible. The study was approved by the ALSPAC Ethics and Law committee, and the Cambridge Human Biology Research Ethics Committee. The participants of the primary MWAS on SCDC were 701 children who completed the SCDC at age 8, and for whom epigenetic data were available (341 males and 360 females). We conducted a secondary MWAS of pragmatic communication in 666 children. Pragmatic communication was measured using the Children’s Communication Checklist^29^ (CCC) at age 9 (323 males and 340 females). In addition, we conducted a GWAS of SCDC scores in a sample of 5,628 8-year olds from ALSPAC, details of which are provided below.

### Phenotypic measures

The SCDC is a 12-item questionnaire that measures difficulties in verbal and nonverbal communication and social interaction including reciprocal interaction^25^. Scores range from 0 to 24 with high scores reflecting difficulties in social interaction and communication. The SCDC has good psychometric properties - internal consistency of 0.93 and test-retest reliability of 0.81^25^. We used mother-reported SCDC scores on children aged 8. The mean of SCDC scores in our samples was 14.65 (standard deviation = 3.44). Previous research has shown that the SCDC is stable over time and scores at different ages are genetically correlated^26,30^. SNP heritability for SCDC measured at various time points is highest in childhood (at the age of 91 months) and in later adolescence (17 years)^26,30^. We focussed on SCDC scores at 91 months as the sample size was the largest, has highest genetic correlation with autism^26^, and the exposure to environmental factors is limited at 91 months compared to other time points.

A second measure that we used in this study, is the 53-item parent-completed CCC which measures pragmatic communication^29^. The CCC and subscales has moderate to high twin heritability (0.53 < h^2^_twin_ < 1)^31^, and moderate SNP heritability (h^2^_SNP_ = 0.18)^32^. There is a negative correlation between the CCC and the SCDC^33^. The mean of the CCC in the sample of 666 children was 151.83 (standard deviation = 6.77), with scores ranging from 111 to 162. To make the analysis comparable with the SCDC (which measures difficulties rather than ability) we reverse scored the CCC so that higher scores measure difficulties in pragmatic communication.

### Cord blood DNA Methylation, quality control and normalization

DNA was extracted from cord blood drawn from the umbilical cord upon delivery. Following extraction, DNA was bisulfite-converted using the Zymo EZ-DNA MethylationTM kit (Zymo, Irvine, CA) then genome-wide methylation status of over 485 000 CpG sites was measured using the Illumina HumanMethylation450 BeadChip array according to the standard protocol. The arrays were scanned using an Illumina iScan and initial quality review was assessed using GenomeStudio (version 2011.1).

Methylation assays utilize a pair of probes to detect methylation of cytosine at CpG sites. One is used to detect methylated loci (M) and the other is used to detect unmethylated CpG islands (U). The level of methylation at a particular locus is then estimated based on the ratio of signals from M to U, called “beta” value. β-values are reported as percentages, ranging from 0 (no cytosine methylation) to 1 (complete cytosine methylation).

### QC and Normalization

In total, there were 1127 cord blood samples including technical replicates. Of these, 241 were from blood spots and 886 were from white cells. 919 of these passed the mother-child genotype-based relatedness quality control. We further removed duplicate samples and participants who were outliers for genetic heterozygosity, genetic ethnicity outliers, and sex mismatch. Samples were further removed if there were low bead numbers, and high detection P-value. This left us with 701 participants who had both epigenetic and phenotypic data.

The data was normalized using functional normalization implemented in the R package meffil (https://github.com/perishky/meffil)34. Functional normalisation is a between-array normalisation method for the Illumina Infinium HumanMethylation450 platform and an extension of quantile normalisation. It removes unwanted technical variation. The normalization procedure was performed to the methylated and unmethylated signal intensities separately. For X and Y chromosomes, males and females were normalised separately using the gender information.

We removed CpG sites whose probe or single-base extension overlaps with a SNP with MAF > 0.01. We further removed cross-reactive probes identified in Chen et al., 2013^35^ as implemented in mefill. In total, 372,662 CpG sites remained after quality control. Cell proportions for CD4 T lymphocytes, CD8 T lymphocytes, B lympocytes, natural killer cells, monocytes, and granulocytes were estimated using the minfi package^36^.

### Methylome-wide association

A methylome-wide association study was run using a two-step regression model. In the first regression, normalized epigenetic probe betas were regressed against technical covariates (slide, sample type, and plates and cell counts). The residuals from this regression were further used as corrected methylation values. In the second regression, SCDC (or CCC) scores were regressed against corrected methylation values with sex and the first two genetic principal components as covariates. Here, we were specifically testing if methylation status measured in cord blood was associated with autistic traits or pragmatic language measured at a later age. Given the highly skewed distribution of the SCDC scores, we used a negative binomial regression, using the MASS package in R. We used a Bonferroni-corrected epigenome-wide significant threshold of 1×10^−7^ to identify significant associations. All analyses were conducted in R version 3.2.

In order to interpret results from the MWAS, we designed a multi-step enrichment strategy including (1) a same-sample, same-tissue overlap and correlation analyses between the SCDC and the CCC; (2) a cross-tissue overlap analysis between the SCDC MWAS and MWAS of autism in peripheral blood and post-mortem brain tissue; (3) enrichment for autism transcriptionally dysregulated genes; and (4) Enrichment of CpG-associated mQTLs in autism and SCDC GWAS. A summary of the study design is provided in **Figure 1**.

**Figure 1:**
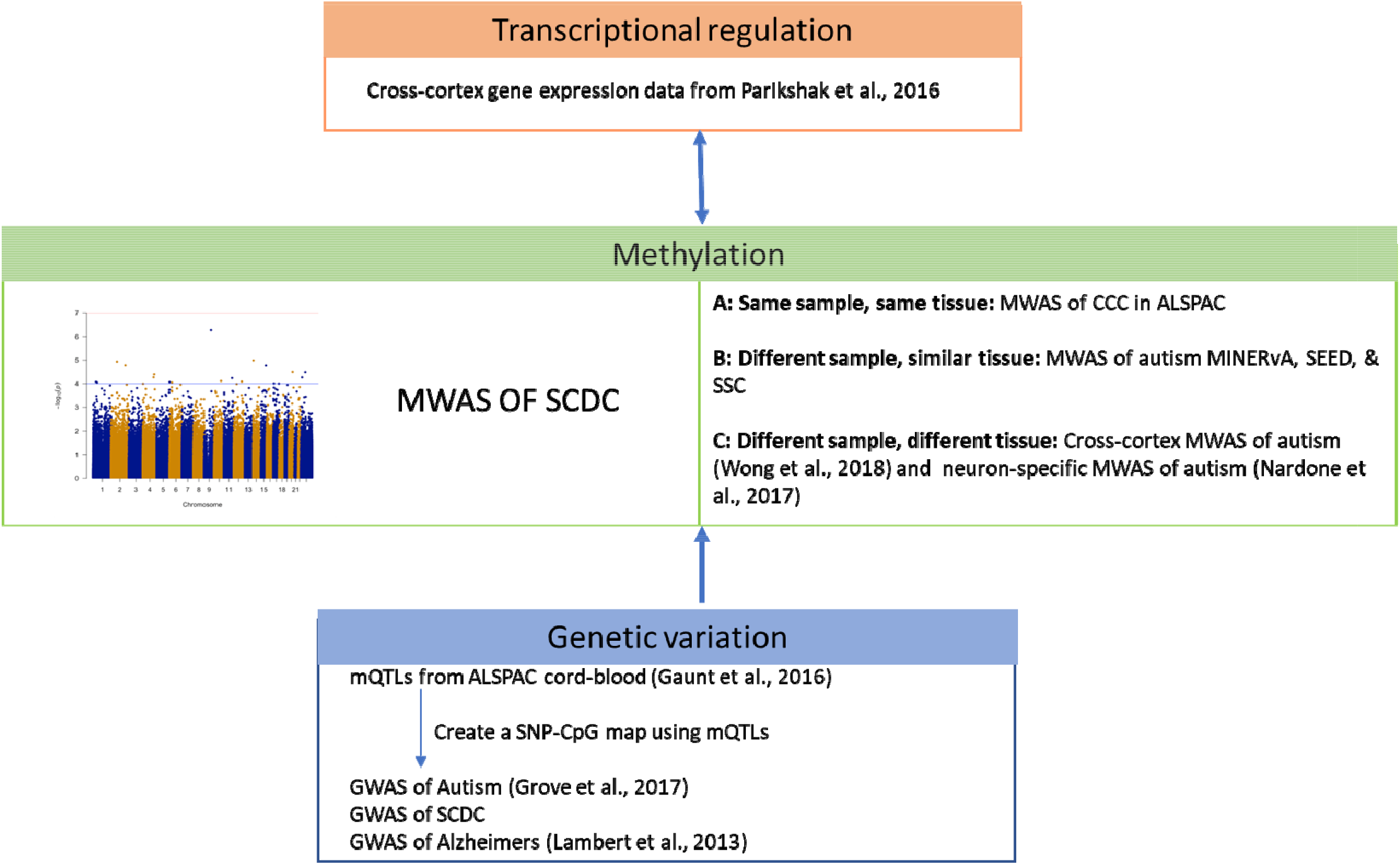
Schematic diagram of the study design

### Peripheral tissue (blood and blood-spot) overlap analysis

We had access to summary MWAS statistics from three peripheral tissue datasets described in detail elsewhere (SEED^19^, Simons Simplex Collection (SSC)^19^, and MINERvA^14^). For all overlap analyses, we conducted two statistical tests. In the first, we included all nominally significant CpGs (P < 0.01) in three peripheral tissue MWAS datasets (SEED, Simons Simplex Collection (SSC), and MINERvA) and tested if these have a shift towards lower P-values in the SCDC MWAS dataset. This tests a larger number of CpGs and is consistent with the idea that each individual CpG contributes minimally to the phenotype suggesting a polymethylomic (or, polyepigenetic) architecture similar to a polygenic architecture of complex traits. In addition, this does not test effect direction as effect direction may vary based on number of factors including tissue source. In the second, we tested for effect direction concordance for CpGs with P < 1×10^−4^ in either of the two datasets being tested, thus, conducting 12 tests in total. We evaluated the significance of the concordance in effect direction using binomial sign test, and corrected for all 12 tests conducted. This restricts the analyses to a relatively small number of CpGs.

### Post-mortem brain tissue overlap analysis

We used two sources of post-mortem brain tissue to investigate overlap with the SCDC MWAS. MWAS in both these datasets were conducted using the Illumina HumanMethylation450 BeadChip making the MWAS comparable to the SCDC MWAS. We used a recent MWAS conducted using tissue from 38 ‘idiopathic’ autistic individuals and 38 non-psychiatric controls^23^. MWAS was conducted using a multi-level linear mixed model with autism status as the independent variable and methylation as the dependent variable, combining samples from the prefrontal cortex and the temporal cortex given the high correlation in methylation values between the two cortical regions. Covariates include age, sex, brain bank and neuronal cell proportions. Further details are provided elsewhere^23^. To further investigate if there was enrichment for neuron-specific methylation signatures, we used MWAS data from FACS-sorted neurons in brain samples from 15 autistic individuals and 16 typical controls^20^. For most individuals tissue was obtained from the BA10 (anterior prefrontal cortex). MWAS was conducted to identify differentially methylated CpGs using a linear model. Further details can be found elsewhere^20^.

For both the datasets, our analysis was similar to the analysis of peripheral tissue MWAS. We investigated effect direction concordance between the two post-mortem brain autism MWAS and the SCDC MWAS for all CpGs with P < 1×10^−4^ in the post-mortem brain MWAS. Additionally, we investigated if CpGs with P < 0.01 in either of the two postmortem brain MWAS had a significant shift towards lower P-values in the SCDC MWAS.

### Enrichment with autism-associated transcriptionally dysregulated genes

For enrichment analyses with transcriptionally dysregulated gene expression data, we used an RNA-sequencing study of 167 post-mortem cortical samples with n = 85 with a diagnosis of autism and n = 82 from nonpsychiatric controls. Samples were from BA9 (prefrontal cortex), or BA41/42 (temporal cortex)^38^. Significantly dysregulated genes had a Benjamini-Hochberg adjusted FDR < 0.05. We conducted enrichment analyses using a onesided Wilcoxon rank-sum test. We first mapped the CpGs to genes using the CpG to gene annotation for the Illumina 450k methylation array using the *IlluminaHumanMethylation450k.db* package in R (http://www.bioconductor.org/packages/release/data/annotation/html/IlluminaHumanMethylation450k.db.html). We restricted our analysis only to CpGs that were mapped onto the genes tested for differential expression in the post-mortem brain dataset^38^. We then compared the distribution of the SCDC P-values for CpGs mapped to significantly differentially dysregulated genes vs the other genes.

### Enrichment of CpG-associated mQTLs in autism and SCDC GWAS

We investigated if mQTLs of CpGs below 4 P-value thresholds in the SCDC MWAS (P_SCDC_) had lower P-values compared to other mQTLs in the GWAS (P_GWAS_) of (1) autism, (2) SCDC, and, (3) as a negative control, Alzheimer. We hypothesized that the mQTLs of CpGs below P_SCDC_ will have significantly lower P_GWAS_ in comparison with remaining mQTLS. To map CpGs to mQTLS we used mQTL maps generated by the ARIES cohort in cord blood, restricting our analysis to only significant mQTLS identified after FDR correction^39^. All mQTLs had a minor allele frequency > 1%. For each CpG-mQTL pair, we restricted our analysis to only those CpG-mQTL pairs investigated in both the SCDC MWAS and the GWAS of interest. In other words, the CpGs must have been investigated in the SCDC MWAS and the paired mQTL of the CpG must have been investigated in the GWAS of interest. As none of the CpGs meet the strict p-value threshold, we had to use several thresholds at different levels of stringency. To control the signal-to-noise ratio in the context of an MWAS, we have considered four empirical P_SCDC_ thresholds: 0.05, 0.01, 0.005, and 0.001. Enrichment was conducted using permutation testing, where we defined the 10,000 null sets. We identified 3 potential factors that may influence this analysis: (1) the Linkage Disequilibrium (LD) structure of mQTLs, (2) the number of mQTLs mapped onto a CpG, (3) the number of CpGs a single mQTL is mapped onto. To address LD, first, we clumped the list of mQTLs using an r^2^ of 0.6 and distance of 1000 kb, to ensure that linkage disequilibrium among these mQTLs does not confound the outcome. In this clumped list of mQTLs, the majority were mapped to only one mQTL. Second, to account for the number of mQTLs mapped onto CpGs, we binned the CpGs into 6 groups based on the number of SNPs they mapped onto (1 – 5, 6 – 10, 11 – 15, 15 – 20, 20 – 25 and above 25), and conducted enrichment analysis so that every mQTL in the null set matched the original mQTL based on CpG bins. Third, one single mQTL may map onto multiple CpGs, resulting in non-unique CpG-mQTL pairs with P_SCDC_ < threshold, and P_SCDC_ > threshold. We retained unique CpG-mQTL pairs in each list before conducting permutation-based enrichment analysis. Finally, to account for multiple testing, as we tested across four non-independent P-value thresholds, the empirical P-values were corrected for the 4 tests using Benjamini-Hochberg FDR correction. Empirical P-values were significant at FDR < 0.05.

We validated the results identified in the Autism GWAS using a GWAS of log-transformed SCDC scores in ALSPAC (details below). As a negative control, we used GWAS data for Alzheimer’s (Phase I), downloaded from IGAP (http://web.pasteur-lille.fr/en/recherche/u744/igap/igap_download.php)^40^, and tested for enrichment using an identical procedure as mentioned above. The Alzheimer’s GWAS (Phase I, for which genome-wide summary data is available) consists of 17,008 cases and 37,154 controls, and identified 14 significant GWAS loci. Whilst both autism and Alzheimer’s are neuropsychiatric conditions, the genetic correlation between the two conditions is nonsignificant (r_g_ = 0.04±0.10; P = 0.102), suggesting minimal shared genetics. The number of cases and controls used in the two studies (Phase 1 for the Alzheimer’s GWAS) are comparable, providing approximately similar statistical power (Mean chi-square: Alzheimer’s = 1.114, Autism = 1.2). Further, they are distinct in that autism is a neurodevelopmental condition diagnosable at childhood, while Alzheimer’s is largely diagnosed in individuals who are 65 or older.

### GWAS of SCDC scores

We conducted a log-transformed genome-wide association study of SCDC scores at age 8 in the ALSPAC data. Note that log-transformed phenotype models are computationally more efficient for high dimensional GWAS data than negative binomial models used in the MWAS. Participants were genotyped using the Illumina^®^ HumanHap550 quad chip by Sample Logistics and Genotyping Facilities at Wellcome Sanger Institute and LabCorp (Laboratory Corportation of America) using support from 23andMe. We restricted our analysis only to individuals of European descent. This was identified using multidimensional scaling analysis and compared with Hapmap II (release 22)^41^. We excluded individuals with gender mismatches, high missingness (> 3%), and disproportionate heterozygosity, and if cryptic relatedness, identified using identity by descent, was greater than 0.1. We removed SNPs with greater than 5% missingness, those that violated Hardy-Weinberg equilibrium (P < 1×10^−6^), and those with a minor-allele frequency less than 1%. This resulted in a total of 526,688 genotyped SNPs. Haplotypes were estimated using data from mothers and children using ShapeIT (v2.r644)^42^. Imputation was performed using Impute2 V2.2.2 against the 1000 genomes reference panel (Phase 1, Version 3)^43^. Imputed SNPs were excluded from all further analyses if they had a minor allele frequency < 1% and info < 0.8. After quality control, there were 8,282,911 genotyped and imputed SNPs that were included in subsequent analyses. GWAS analysis was conducted for mother-reported SCDC scores at age 8 that was log-transformed given the highly skewed distribution. Linear regression was conducted in Plink v1.9^44^ that converted allele dosages into hard calls. We included the first two ancestry principal components and sex as covariates in the regression model.

As reported previously^5,26,30^, the SNP heritability as quantified using LDSC^45,46^ was h^2^ = 0.12 ± 0.05. The LDSR intercept (0.99) suggested that there was no inflation in GWAS estimates due to population stratification. The λ_GC_ was 1.013. We replicated the previously identified genetic correlation (constrained intercept)^5^ with autism using our SCDC GWAS (PGC-autism: r_g_ = 0.46 ±0.20, P = 0.019; iPSYCH-autism: r_g_ = 0.45±0.18, P = 0.01).

### Data, software, and script availability

a. MWAS summary statistics:

- The summary statistics for the MWAS (SCDC and CCC) can be downloaded from here: https://www.dropbox.com/sh/8za5xspmbjydpst/AAA_ZGmMLOE8Ql7egi5Mcu8Ha?dl=0.
- Summary statistics for the SEED and the SSC MWAS can be obtained from here: https://molecularautism.biomedcentral.com/articles/10.1186/s13229-018-0224-6.
- Summary statistics for the MINERvA cohort can be obtained by contacting Jonas Bybjerg-Grauholm.
b. GWAS summary statistics:

- The summary statistics for the autism GWAS (iPSYCH) can be downloaded from http://www.med.unc.edu/pgc/results-and-downloads (iPSYCH-PGC GWAS-2017).
- The Alzheimer’s GWAS can be downloaded from http://web.pasteur-lille.fr/en/recherche/u744/igap/igap_download.php.
- The summary statistics for the SCDC GWAS can be obtained from https://www.dropbox.com/sh/8za5xspmbjydpst/AAA_ZGmMLOE8Ql7egi5Mcu8Ha?dl=0.
c. Scripts for running the two regression models for the MWAS and running the enrichment analyses with the mQTL data are available here: https://github.com/autism-research-centre/MWAS_autistictraits
d. mQTL data used in this (coord blood) is a part of the ARIES cohort, and can be downloaded here: http://www.mqtldb.org/
e. We used the following softwares/packages: Plink (http://zzz.bwh.harvard.edu/plink/); IlluminaHumanMethylation450k.db (http://www.bioconductor.org/packages/release/data/annotation/html/IlluminaHumanMethylation450k.db.html); MASS (https://cran.r-project.org/web/packages/MASS/index.html); LDSC (https://github.com/bulik/ldsc/wiki/Heritability-and-Genetic-Correlation).

## Results

### Methylome-wide association study of the SCDC scores

To identify if there are any significantly associated CpG sites with SCDC, we conducted a methylome-wide association study. Methylome-wide association analysis did not identify any significant loci after Bonferroni correction (P<1×10^−7^). The top CpG site was cg14379490, on chromosome 9 (MWAS Beta = −1.78±0.35, P = 5.34×10^−*7*^). This CpG site is an ‘Open Sea’ CpG site, whose closest gene is *FAM120A*, which encodes a scaffold protein that is expressed in a wide number of human tissues. We identified 19 CpG sites with suggestive P-values (P < 1×10^−4^) (**Supplementary Table 1**). The QQplot and the Manhattan plot are provided in **Figure 2**. We did not find any evidence for inflation in P-values (λ = 0.88), possibly because of the relatively small sample size and the regression model used.

**Figure 2:**
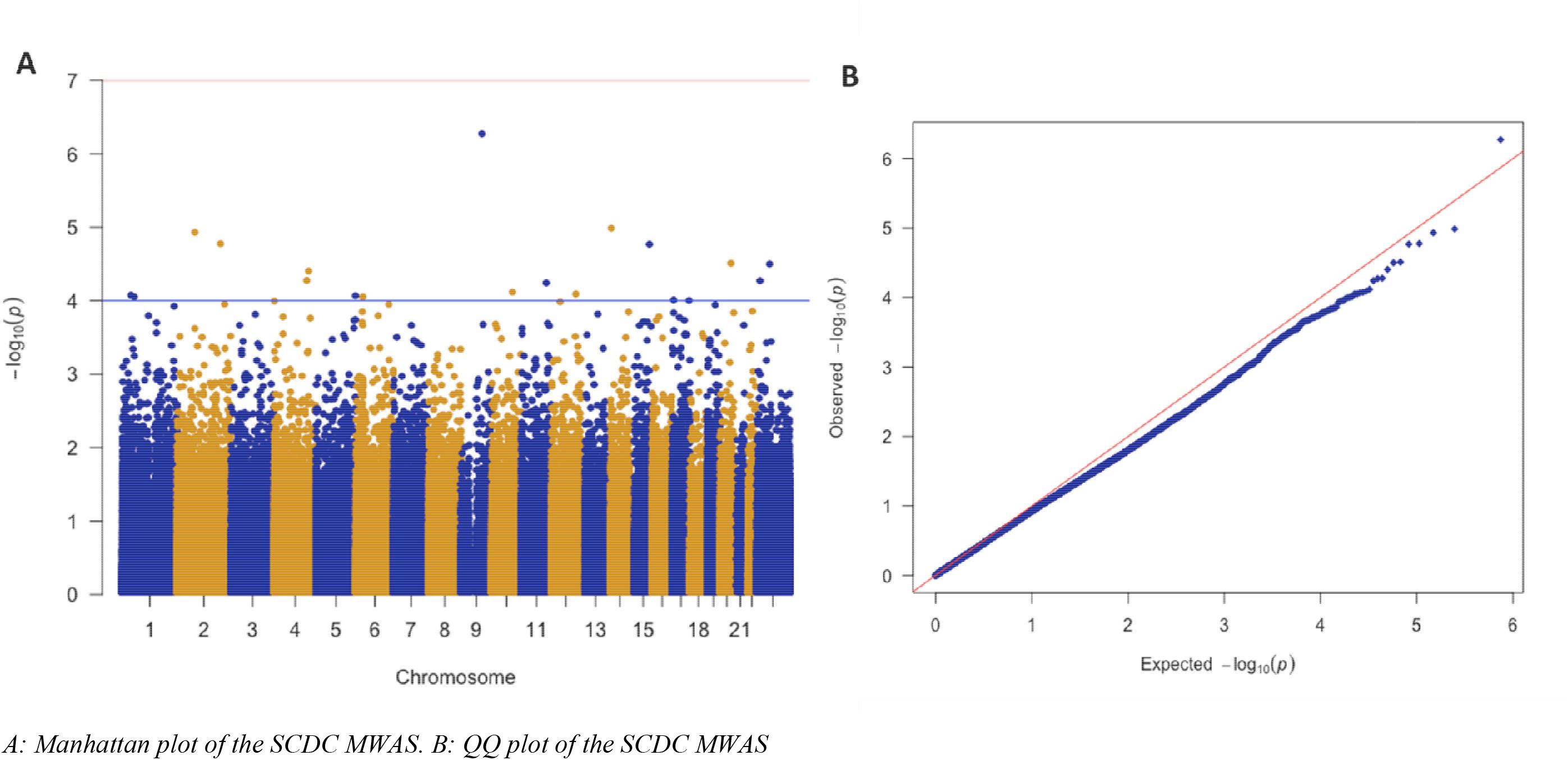
Manhattan plot and QQ plot for the SCDC MWAS

To confirm that the signals were biologically meaningful, we additionally conducted a MWAS of the Children’s Communication Checklist (CCC) reversed scored to identify difficulties in communication (**Methods**). The most significant CpG was cg13711424 (MWAS Beta = −3.73±0.71, P = 1.79×10^−7^). The Manhattan plot and QQ plot are included in **Supplementary Figure 1**. Of the 19 SCDC-associated CpGs of suggestive significance (P < 1×10^−4^), the effect was concordant for 18 of them in the CCC MWAS (P = 7.62×10^−5^, binomial sign test). Similarly, of the 32 CpGs with P < 1×10^−4^ in the CCC MWAS, 28 had concordant effect direction in the SCDC MWAS (P = 1.93×10^−5^, binomial sign test). Scores on the CCC and the SCDC were phenotypically correlated (r = 0.39, 95% CI = 0.32 - 0.45, P < 2.2×10^−16^) in the participants who were included in the MWAS (n = 666), and both questionnaires measure difficulties in pragmatic communication. Given that we were testing correlated phenotypes in the same cohort using methylation signatures form the same tissues, we hypothesized that effect should be positively correlated for the two MWAS. The Z-scores for the MWAS for the two phenotypes were significantly correlated (r = 0.157, 95% CI = 0.153 – 0.160, P < 2.2×10^−16^), which increased if we considered the CpGs with P < 0.01 in either one of the phenotypes (P_SCDC_ < 0.01: r = 0.40, 95% CI = 0.36 – 0.43, P < 2.2×10^−16^, P_CCC_ < 0.01: r = 0.40, 95%CI = 0.37 − 0.42, P < 2.2×10^−16^).

### Enrichment analyses with peripheral blood methylation signatures

To investigate if there is an overlap between the SCDC MWAS with MWAS of autism conducted in peripheral tissues, we conducted effect direction concordance analysis with three autism datasets (MINERvA, SEED, and SSC, **Methods**). For all of them, we first investigated concordance of effect direction of all CpG sites with P < 1×10^−4^. In contrast to the findings with the CCC, we did not identify a significant concordance in effect direction between the SCDC MWAS and any of the other three autism MWAS datasets. Indeed, none of the three MWAS datasets had significant concordance in effect direction for the suggestive CpGs in each analysis (**Table 1**).

**Table 1:** Sign concordance of the SCDC MWAS and the three peripheral tissue MWAS at top loci (P < 1×10^−4^)

|  |  | Testing dataset |  |  |  | Number of CpGs in the discovery dataset |
| --- | --- | --- | --- | --- | --- | --- |
|  |  | ALSPAC | MINERVA | SEED | SSC |  |
| **Discovery dataset** | ALSPAC | **NA** | **10** | **11** | **4** | 17 |
|  | MINERVA | **14** | **NA** | **17** | **19** | 29 |
|  | SEED | **19** | **21** | **NA** | **18** | 37 |
|  | SSC | **19** | **21** | **23** | **NA** | 47 |

*The table above provides a test of effect direction concordance. CpGs with $P < 1 \times 10^{-4}$ in the discovery dataset were tested for concordance in effect direction in the testing dataset. The numbers in the cells (in bold) the provides the total number of CpGs with conocondant effect direction in the testing dataset. The Number of CpGs in the discovery dataset provides the total number of CpGs in the discovery dataset with $P < 1 \times 10^{-4}$ . None of the results were significant (binomial sign test) after correcting for the multiple tests conducted.*

Given that there was limited evidence for concordance in effect direction between the datasets, we next tested if nominally significant CpGs (P < 0.01) in the three autism MWAS have a shift towards lower P-values in the SCDC MWAS using a one-sided Wilcoxon-rank sum test. This tests more CpGs than an effect direction concordance test does, and is agnostic to effect direction which may be discordant in different peripheral tissues measured at different developmental stages. After Bonferroni correction (alpha = 0.016), we did not identify a significant shift towards lower P-values for the nominally significant CpGs from any of the three datasets (SEED: P = 0.02; SSC: P = 0.48; MINERvA: P = 0.91), though we note a nominally significant shift in the SEED dataset.

### Enrichment analyses with autism postmortem methylation signatures

Methylation signatures in postmortem brain tissues are more relevant to neurodevelopmental phenotypes than methylation signatures in peripheral tissue. Considering this, we investigated if there is an enrichment between the SCDC MWAS and MWAS of the postmortem autism brain. Using data from the latest post-mortem brain study^23^, we investigated concordance in effect direction between all CpG probes with P < 1×10^−4^ from the cross-cortex analysis in the SCDC MWAS. 171 out of 293 CpGs had a concordant effect direction in the two datasets (P = 0.004). At a more stringent P-value threshold of P < 1×10^−5^, 88 of the 133 probes had concordant effect directions in the two datasets (P = 2.4×10^−4^, binomial sign test). In contrast, Wilcoxon rank-sum test of all CpGs with P < 0.01 in the postmortem MWAS did not identify a significant shift towards lower P-values (P = 0.99, one-tailed Wilcoxon rank-sum test). We next tested if we could validate the concordance in effect direction in a different dataset. A previous study has investigated differential methylation in post-mortem neurons from the frontal lobe (identified using FACS sorting) in autism^20^. First, testing effect direction concordance, 44 of the 87 CpGs with P < 1×10^−4^ had concordant effect direction in the two datasets (P = 1, binomial sign test). However, we identified a significant shift towards lower P-values (P = 9.3 x 10^−3^, one-tailed Wilcoxon rank-sum test) of all CpGs with P < 0.01 in the SCDC MWAS.

### Enrichment with autism dysregulated genes

A few studies have identified consistent sets of dysregulated genes in autism, and co-expression modules enriched^28–31^. Previous studies have identified a significant enrichment for differentially methylated CpGs and genes that are transcriptionally dysregulated in the post-mortem cortex in autism^14^. We investigated if CpGs mapped to transcriptionally dysregulated genes in the autism postmortem cortex^38^ and associated co-expression had a shift towards lower P-values in the SCDC MWAS when compared to the other genes. We identified a significant shift towards lower P-values for the transcriptionally dysregulated genes (One-sided Wilcoxon rank-sum test, P = 6.22×10^−5^), but did not identify a significant enrichment for any of the modules (M4: P = 0.58, M9: P = 0.59, M16: P = 0.042, M10: P = 0.31, M20: P = 0.42, M19: P = 0.105).

### Genetic influences in SCDC methylation patterns

We next investigated if the methylation signatures associated with SCDC scores are enriched for GWAS signals for autism. DNA methylation is under cis and, to a smaller extent, trans genetic control. We identified mQTLS associated with SCDC CpG probes below 4 P-value thresholds (P_SCDC_, Methods), and compared the distribution of P-value of the mQTLS in the autism GWAS against the P-value distributions of mQTLs above the P_SCDC_ (**Methods**). After multiple testing correction, mQTLS of CpGs with P_SCDC_ = 0.01, and 0.005 has significantly lower P-values in the autism GWAS dataset (P_SCDC_ 0.01: P__FDRcorrected_ = 5×10^−4^, P_SCDC_ 0.005, P__FDRcorrected_ = 4.75×10^−3^) (**Table 2, Figure 2**). We validated this enrichment in a GWAS of SCDC, which is genetically correlated with autism. We identified an enrichment at P_SCDC_ 0.005 (P__FDRcorrected_ = 0.046) and at PSCDC 0.001 (P__FDRcorrected_ = 0.046). In contrast, we did not identify an enrichment mQTLs in the Alzheimer’s GWAS (**Table 2, Figure 3**).

**Table 2:** Results of the enrichment analysis of the top CpGs

| **CpG P-value threshold** | **GWAS** | **P** | **Mean difference** | **FDR P** |
| --- | --- | --- | --- | --- |
| **0.05:** | Autism | 0.54 | 0.004 | 0.54 |
| **0.01:** | Autism | $1.0 \times 10^{-4}$ | 0.039 | $5 \times 10^{-4}$ |
| **0.005** | Autism | $1.9 \times 10^{-3}$ | 0.031 | $4.75 \times 10^{-3}$ |
| **0.001:** | Autism | 0.040 | 0.037 | 0.063 |
| **0.05:** | SCDC | 0.735 | 0.007 | 0.073 |
| **0.01:** | SCDC | 0.301 | 0.015 | 0.40 |
| **0.005** | SCDC | 0.022 | 0.038 | 0.046 |
| **0.001:** | SCDC | 0.023 | 0.065 | 0.046 |
| **0.05:** | Alzheimer's | 0.343 | 0.008 | 0.853 |
| **0.01:** | Alzheimer's | 0.710 | 0.003 | 0.853 |
| **0.005** | Alzheimer's | 0.853 | -0.003 | 0.853 |
| **0.001:** | Alzheimer's | 0.793 | -0.009 | 0.853 |

*The table provides the results of the enrichment analyses for the top loci. Mean difference is the difference in average GWAS P-values for the mQTLs mapped to CpGs below the threshold in the MWAS from the average GWAS P-values for the mQTLs mapped to CPGs in the MWAS. A positive difference suggests and enrichment.*

**Figure 3:**
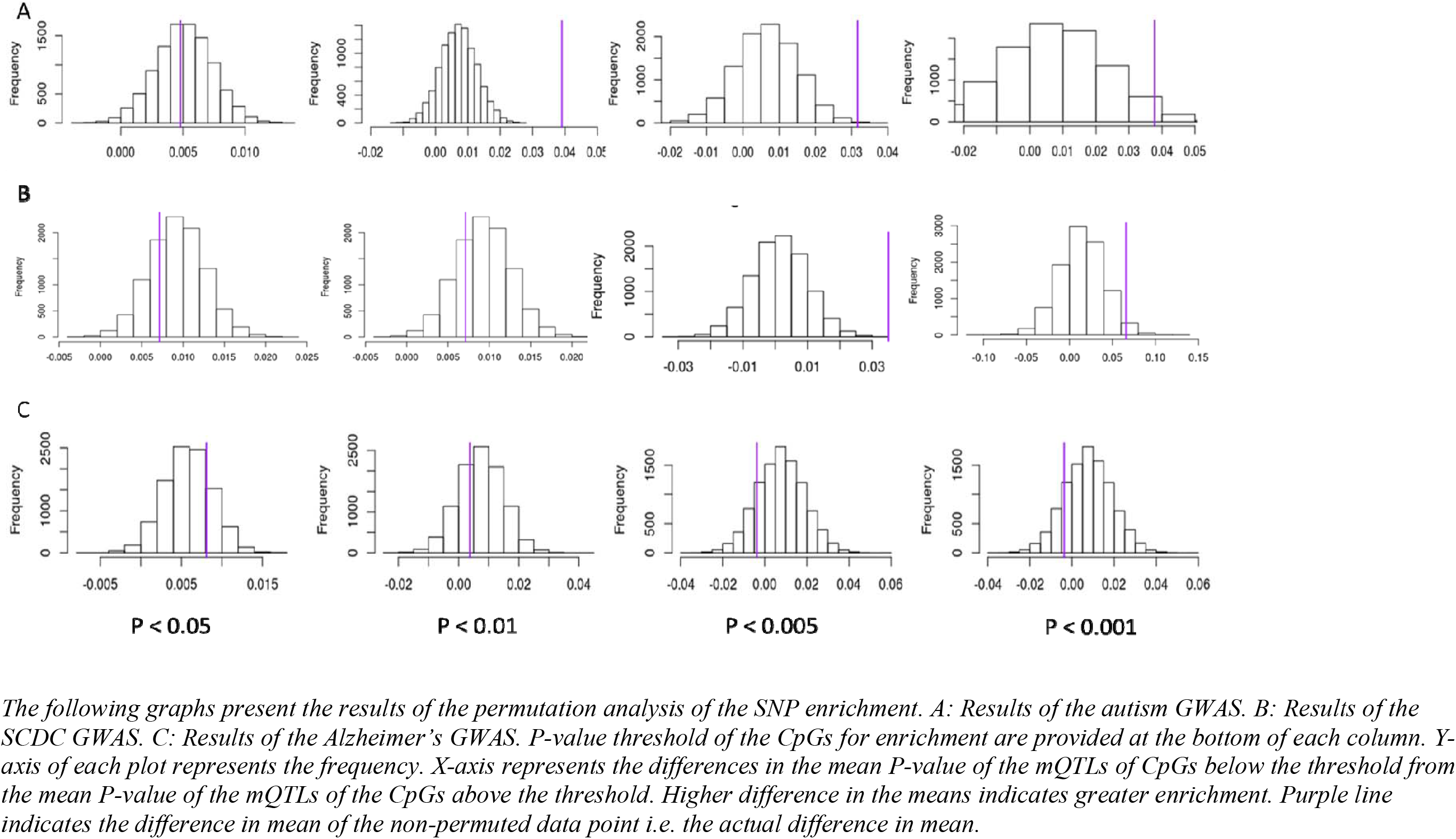
Permutation histogram of SNP-enrichment in top CpGs for three GWAS

## Discussion

This study investigates the shared biology of autism and autistic traits by integrating genetic, methylation data, and post-mortem gene expression data. We first investigated the validity of considering autistic traits for methylation studies. Considering autistic traits over a case-control design is useful in that: 1. It captures greater variance across the underlying liability spectrum; 2. It can be used to increase sample sizes by phenotyping individuals for whom methylation data is available; and 3. It can be used to link methylation signatures from tissues collected in early life to the phenotype, as this can be more difficult for autism.

We conducted a prospective MWAS of autistic traits (SCDC) by measuring methylation signatures in the cord blood and linking it to autistic traits measured later in life. Whilst we did not identify a significant CpG association with autistic traits after multiple testing correction, we were able to confirm that this analysis produced biologically meaningful signal by identifying significant correlation with an MWAS of a similar childhood phenotype (pragmatic language phenotype) measured in the same cohort. Notably, the correlation in methylation values mirrored the phenotypic correlation, providing confidence in our results.

Despite this, however, we did not identify a significant overlap between the MWAS of SCDC and MWAS of autism conducted using peripheral tissues^14,19^. We note several differences in the SCDC MWAS analysis and the three MWAS. Of primary importance is the statistical model used in the analysis. Whilst we were interested in investigating if methylation signatures from cord blood were associated with SCDC scores measured in later life, all three peripheral tissue MWAS investigated if autism diagnosis was associated with differential methylation. Thus, in our analysis, methylation was an independent and continuous variable, whereas in the three MWAS, it was a dependent and discrete variable. Second, there are remarkable differences in tissue source, age at which methylation was measured, and confounding variables included in the analyses (for instance we included genetic principle components as covariates). Interestingly, none of the three autism MWAS showed significant overlap with each other as investigated using a sign-concordance test of the most significant CpGs. It is critical to investigate this observed lack of concordance

In contrast to the results from the peripheral tissues, we observed some degree of overlap between MWAS conducted in post-mortem brain tissues^20,23^ and the SCDC MWAS. First, we found a significant sign concordance in CpGs identified in the largest cross-cortex MWAS of autism using postmortem tissue samples. However, we did not identify an enrichment using a Wilcoxon-rank sum test of P-values. In contrast, using a neuron-specific MWAS generated using different post-mortem tissue, we identified a significant overlap using a Wilcoxon-rank sum test of P-values but not a sign-concordance in this dataset. Additionally, using an RNA sequencing dataset of autism and neurotypical postmortem brains^38^, we again identified a significant enrichment for transcriptionally dysregulated genes using a Wilcoxon-rank sum test. Overall, we are unable to strongly suggest that there is a significant overlap between the SCDC MWAS and the MWAS of autism in either postmortem or peripheral blood tissues. This is likely due to multiple factors as outlined earlier. In addition, measuring methylation in peripheral tissue, which is not central to a neurodevelopmental condition like autism, is likely to attenuate the signal to noise ratio. Indeed, the post-mortem brain MWAS study^23^ has identified significant CpGs with fewer samples compared with any of the three peripheral tissue MWAS^14,19^. Thus, due to both the increased statistical power and the use of the relevant tissue, the top CpGs in the post-mortem brain MWAS are more likely to be true positives than the top CpGs in the peripheral tissue MWAS.

Given the highly polygenic nature of autism^11^, it is likely that GWAS loci that are not statistically significant in the current GWAS studies may still influence methylation. Thus, our second aim of this study was to investigate if GWAS signals for autism and autistic traits are enriched in the top CpG sites in the autism MWAS by using mQTL mapping. Our results demonstrate an enrichment for mQTLs for CpGs associated with SCDC scores in the GWAS for autism. Crucially, we were able to validate the results in a much smaller GWAS of SCDC scores, but failed to identify an enrichment in a GWAS of Alzheimer’s^40^, which is of comparable statistical power to the GWAS of autism. This enrichment is observed at more stringent P-value thresholds providing confidence in our results. We did not test this in other peripheral tissue MWAS for which we had access to summary statistics given the marked lack of overlap between these and the SCDC MWAS.

Our study does not, however, investigate causality. While methods such as Mendelian randomization can investigate causality^14,47^, this is typically restricted to a few number of loci based on current results of GWAS studies. In addition, we are restricted from using mendelian randomization due to the low statistical power of both the MWAS and the GWAS sets, resulting in the identification of a limited number of statistically significant loci. Two mechanisms may explain the overlap observed in the current dataset. The first is causal in nature, wherein, genetic loci are likely to influence autism or autistic traits by influencing methylation levels of CpG sites. This can influence gene expression levels. The second is horizontal pleiotropy, where genetic loci are associated with autism or autistic traits, and separately, also influence methylation levels of CpG sites. This study cannot tease these two mechanisms apart.

Three caveats must be borne in mind whilst interpreting the results of this analysis. First, the current array-based method interrogates only a small proportion of all the CpG sites in the genome. Thus, significant loci associated with autistic traits may lie outside of the regions interrogated. Second, due to the nature of the assay, the methylation values may also capture hydroxymethylation. We cannot exclude the possibility of signal attenuation due to assaying both hydroxymethylation and methylation in the current study, and the correlation between hydroxymethylation between blood and brain is low^48^. Third, whilst there is a modest but significant genetic and phenotypic correlation between autism and the SCDC, the SCDC only measures social aspects of autism and is not correlated with the non-social aspects of autism.

Our study demonstrates a degree of overlap between autism and autistic traits, but we are limited in making further conclusions. Two factors – sample size and heterogeneity between the various samples limit our understanding of methylation in autism. We identify an enrichment for autism and autistic traits GWAS signals in the top CpG loci for autistic trait, but these must be replicated in independent MWAS of autistic traits in cord blood.

## Supporting information

## Acknowledgements

We are grateful to all the families who took part in this study, the midwives for their help in recruiting them, and the whole ALSPAC team, which includes interviewers, computer and laboratory technicians, clerical workers, research scientists, volunteers, managers, receptionists, and nurses. This study was funded by grants from the Medical Research Council, the Wellcome Trust, the Autism Research Trust, and the Templeton World Charity Foundation. The research was conducted in association with the National Institute for Health Research (NIHR) Collaboration for Leadership in Applied Health Research and Care East of England at Cambridgeshire and Peterborough NHS Foundation Trust. The views expressed are those of the author(s) and not necessarily those of the NHS, the NIHR or the Department of Health. We also thank the NIHR Cambridge Biomedical Research Centre for support. The UK Medical Research Council and Wellcome (grant ref: 102215/2/13/2) and the University of Bristol provide core support for ALSPAC. GWAS data was generated by Sample Logistics and Genotyping Facilities at Wellcome Sanger Institute and LabCorp (Laboratory Corporation of America) using support from 23andMe. This publication is the work of the authors who will serve as guarantors for the content of this paper. This study was supported by grant HD073978 from the Eunice Kennedy Shriver National Institute of Child Health and Human Development, National Institute of Environmental Health Sciences, and National Institute of Neurological Disorders and Stroke; and by the Beatrice and Samuel A. Seaver Foundation. We acknowledge iPSYCH and The Lundbeck Foundation for providing samples and funding for the MINERvA dataset. The iPSYCH (The Lundbeck Foundation Initiative for Integrative Psychiatric Research) team acknowledges funding from The Lundbeck Foundation (grant numbers R102-A9118 and R155–2014-1724), the Stanley Medical Research Institute, the European Research Council (project number 294838), the Novo Nordisk Foundation for supporting the Danish National Biobank resource, and grants from Aarhus and Copenhagen Universities and University Hospitals, including support to the iSEQ Center, the GenomeDK HPC facility, and the CIRRAU Center. This research has been conducted using the Danish National Biobank resource, supported by the Novo Nordisk Foundation. The SEED study was supported by Centers for Disease Control and Prevention (CDC) Cooperative Agreements announced under the RFAs 01086, 02199, DD11–002, DD06–003, DD04–001, and DD09–002 and the SEED DNA methylation measurements were supported by Autism Speaks Award #7659 to MDF. The SSC was supported by Simons Foundation (SFARI) award and NIH grant MH089606. The project leading to this application has received funding from the Innovative Medicines Initiative 2 Joint Undertaking (JU) under grant agreement No 777394. The JU receives support from the European Union’s Horizon 2020 research and innovation programme and EFPIA and AUTISM SPEAKS, Autistica, SFARI.

## References

1 American Psychiatric Association. Diagnostic and statistical manual of mental disorders (5th ed.). 2013.

2 Lai M-C, Lombardo M V., Baron-Cohen S. Autism. Lancet 2013. doi:10.1016/S0140-6736(13)61539-1.

3 Baron-Cohen S, Wheelwright SJ, Skinner R, Martin J, Clubley E. The autism-spectrum quotient (AQ): evidence from Asperger syndrome/high-functioning autism, males and females, scientists and mathematicians. J Autism Dev Disord 2001; 31: 5–17.

4 Robinson EB, Koenen KC, McCormick MC, Munir K, Hallett V, Happé F et al. Evidence that autistic traits show the same etiology in the general population and at the quantitative extremes (5%, 2.5%, and 1%). Arch Gen Psychiatry 2011; 68: 1113–21.

5 Robinson EB, St Pourcain B, Anttila V, Kosmicki JA, Bulik-Sullivan BK, Grove J et al. Genetic risk for autism spectrum disorders and neuropsychiatric variation in the general population. Nat Genet 2016; 48: 552–5.

6 Colvert E, Tick B, McEwen F, Stewart C, Curran SR, Woodhouse E et al. Heritability of autism spectrum disorder in a UK population-based twin sample. JAMA Psychiatry 2015; 72: 415–23.

7 Tick B, Bolton PF, Happé F, Rutter M, Rijsdijk F. Heritability of autism spectrum disorders: A meta-analysis of twin studies. J Child Psychol Psychiatry Allied Discip 2016; 57: 585–595.

8 Sandin S, Lichtenstein P, Kuja-Halkola R, Hultman C, Larsson H, Reichenberg A. The Heritability of Autism Spectrum Disorder. JAMA 2017; 318: 1182.

9 Hoekstra RA, Bartels M, Verweij CJH, Boomsma DI, PE S, DI B. Heritability of autistic traits in the general population. Arch Pediatr Adolesc Med 2007; 161: 372–7.

10 Sanders SJ, He X, Willsey AJ, Devlin B, Roeder K, State MW et al. Insights into Autism Spectrum Disorder Genomic Architecture and Biology from 71 Risk Loci Article Insights into Autism Spectrum Disorder Genomic Architecture and Biology from 71 Risk Loci. Neuron 2015; 87: 1215–1233.

11 Grove J, Ripke S, Als TD, Mattheisen M, Walters R, Won H et al. Common risk variants identified in autism spectrum disorder. bioRxiv 2017;: 224774.

12 Gaugler T, Klei LL, Sanders SJ, Bodea CA, Goldberg AP, Lee AB et al. Most genetic risk for autism resides with common variation. Nat Genet 2014; 46: 881–5.

13 The Autism Spectrum Disorders Working Group of The Psychiatric Genomics Consortium. Meta-analysis of GWAS of over 16,000 individuals with autism spectrum disorder highlights a novel locus at 10q24.32 and a significant overlap with schizophrenia. Mol Autism 2017; 8: 21.

14 Hannon E, Schendel D, Ladd-Acosta C, Grove J, Hansen CS, Andrews S V. et al. Elevated polygenic burden for autism is associated with differential DNA methylation at birth. Genome Med 2018; 10: 19.

15 Hannon E, Knox O, Sugden K, Burrage J, Wong CCY, Belsky DW et al. Characterizing genetic and environmental influences on variable DNA methylation using monozygotic and dizygotic twins. PLOS Genet 2018; 14: e1007544.

16 Gordon L, Joo JE, Powell JE, Ollikainen M, Novakovic B, Li X et al. Neonatal DNA methylation profile in human twins is specified by a complex interplay between intrauterine environmental and genetic factors, subject to tissue-specific influence. Genome Res 2012; 22: 1395–1406.

17 van Dongen J, Nivard MG, Willemsen G, Hottenga J-J, Helmer Q, Dolan C V. et al. Genetic and environmental influences interact with age and sex in shaping the human methylome. Nat Commun 2016; 7: 11115.

18 Wong CCY, Meaburn EL, Ronald A, Price TS, Jeffries AR, Schalkwyk LC et al. Methylomic analysis of monozygotic twins discordant for autism spectrum disorder and related behavioural traits. Mol Psychiatry 2014; 19: 495–503.

19 Andrews S V., Sheppard B, Windham GC, Schieve LA, Schendel DE, Croen LA et al. Case-control meta-analysis of blood DNA methylation and autism spectrum disorder. Mol Autism 2018; 9: 40.

20 Nardone S, Sams DS, Zito A, Reuveni E, Elliott E. Dysregulation of Cortical Neuron DNA Methylation Profile in Autism Spectrum Disorder. Cereb Cortex 2017; 27: 5739–5754.

21 Nardone S, Sams DS, Reuveni E, Getselter D, Oron O, Karpuj M et al. DNA methylation analysis of the autistic brain reveals multiple dysregulated biological pathways. Transl Psychiatry 2014; 4: 1–9.

22 Ladd-Acosta C, Hansen KD, Briem E, Fallin MD, Walter, Kaufmann E et al. Common DNA methylation alterations in multiple brain regions in autism. Mol Psychiatry 2014; 19: 862–871.

23 Wong C, Smith R, Hannon E, Ramaswami G, Parikshak N, Assary E et al. Genome-wide DNA methylation profiling identifies convergent molecular signatures associated with idiopathic and syndromic forms of autism in postmortem human brain tissue. bioRxiv 2018;: 394387.

24 Andrews S V., Ellis SE, Bakulski KM, Sheppard B, Croen LA, Hertz-Picciotto I et al. Cross-tissue integration of genetic and epigenetic data offers insight into autism spectrum disorder. Nat Commun 2017; 8: 1011.

25 Skuse DH, Mandy WPL, Scourfield J. Measuring autistic traits: heritability, reliability and validity of the Social and Communication Disorders Checklist. Br J Psychiatry 2005; 187: 568–572.

26 St Pourcain B, Robinson EB, Anttila V, Sullivan BB, Maller J, Golding J et al. ASD and schizophrenia show distinct developmental profiles in common genetic overlap with population-based social communication difficulties. Mol Psychiatry 2017. doi:10.1038/mp.2016.198.

27 Boyd A, Golding J, Macleod J, Lawlor DA, Fraser A, Henderson J et al. Cohort Profile: The ‘Children of the 90s’—the index offspring of the Avon Longitudinal Study of Parents and Children. Int J Epidemiol 2013; 42: 111–127.

28 Relton CL, Gaunt T, McArdle W, Ho K, Duggirala A, Shihab H et al. Data Resource Profile: Accessible Resource for Integrated Epigenomic Studies (ARIES). Int J Epidemiol 2015; 44: 1181–1190.

29 Bishop D. Development of the Children’s Communication Checklist (CCC): a method for assessing qualitative aspects of communicative impairment in children. J Child Psychol Psychiatry 1998; 39: 879–91.

30 St Pourcain B, Skuse DH, Mandy WP, Wang K, Hakonarson H, Timpson NJ et al. Variability in the common genetic architecture of social-communication spectrum phenotypes during childhood and adolescence. Mol Autism 2014; 5: 18.

31 Bishop DVM, Laws G, Adams C, Norbury CF. High Heritability of Speech and Language Impairments in 6-year-old Twins Demonstrated Using Parent and Teacher Report. Behav Genet 2006; 36: 173–184.

32 St Pourcain B, Whitehouse AJO, Ang WQ, Warrington NM, Glessner JT, Wang K et al. Common variation contributes to the genetic architecture of social communication traits. Mol Autism 2013; 4: 34.

33 Skuse DH, Mandy W, Steer C, Miller LL, Goodman R, Lawrence K et al. Social Communication Competence and Functional Adaptation in a General Population of Children: Preliminary Evidence for Sex-by-Verbal IQ Differential Risk. J Am Acad Child Adolesc Psychiatry 2009; 48: 128–137.

34 Min J, Hemani G, Smith GD, Relton CL, Suderman M. Meffil: efficient normalisation and analysis of very large DNA methylation samples. bioRxiv 2017;: 125963.

35 Chen Y, Lemire M, Choufani S, Butcher DT, Grafodatskaya D, Zanke BW et al. Discovery of cross-reactive probes and polymorphic CpGs in the Illumina Infinium HumanMethylation450 microarray. Epigenetics 2013; 8: 203–209.

36 Aryee MJ, Jaffe AE, Corrada-Bravo H, Ladd-Acosta C, Feinberg AP, Hansen KD et al. Minfi: a flexible and comprehensive Bioconductor package for the analysis of Infinium DNA methylation microarrays. Bioinformatics 2014; 30: 1363–9.

37 Fischbach GD, Lord C. The Simons Simplex Collection: A Resource for Identification of Autism Genetic Risk Factors. Neuron 2010; 68: 192–195.

38 Parikshak NN, Swarup V, Belgard TG, Irimia M, Ramaswami G, Gandal MJ et al. Genome-wide changes in lncRNA, splicing, and regional gene expression patterns in autism. Nature 2016; 540: 423–427.

39 Gaunt TR, Shihab HA, Hemani G, Min JL, Woodward G, Lyttleton O et al. Systematic identification of genetic influences on methylation across the human life course. Genome Biol 2016; 17: 61.

40 Lambert J-C, Ibrahim-Verbaas CA, Harold D, Naj AC, Sims R, Bellenguez C et al. Meta-analysis of 74,046 individuals identifies 11 new susceptibility loci for Alzheimer’s disease. Nat Genet 2013; 45: 1452–1458.

41 Gibbs RA, Belmont JW, Hardenbol P, Willis TD, Yu F, Zhang H et al. The International HapMap Project. Nature 2003; 426: 789–796.

42 Delaneau O, Marchini J, Zagury J-F. A linear complexity phasing method for thousands of genomes. Nat Methods 2011; 9: 179–181.

43 Howie BN, Donnelly P, Marchini J. A flexible and accurate genotype imputation method for the next generation of genome-wide association studies. PLoS Genet 2009; 5: e1000529.

44 Purcell S, Neale B, Todd-Brown K, Thomas L, Ferreira MAR, Bender D et al. PLINK: a tool set for whole-genome association and population-based linkage analyses. Am J Hum Genet 2007; 81: 559–75.

45 Bulik-Sullivan BK, Loh P-R, Finucane HK, Ripke S, Yang J, Patterson N et al. LD Score regression distinguishes confounding from polygenicity in genome-wide association studies. Nat Genet 2015; 47: 291–295.

46 Bulik-Sullivan BK, Finucane HK, Anttila V, Gusev A, Day FR, Loh P-R et al. An atlas of genetic correlations across human diseases and traits. Nat Genet 2015; 47: 1236–41.

47 Hannon E, Weedon M, Bray N, O’Donovan M, Mill J. Pleiotropic Effects of Trait-Associated Genetic Variation on DNA Methylation: Utility for Refining GWAS Loci. Am J Hum Genet 2017; 100: 954–959.

48 Lunnon K, Hannon E, Smith RG, Dempster E, Wong C, Burrage J et al. Variation in 5-hydroxymethylcytosine across human cortex and cerebellum. Genome Biol 2016; 17: 27.

