## Supplementary material for "Integrated genetic and methylomic analyses identify shared biology between autism and autistic traits"

**Supplementary Note**

**iPSYCH-Minerva Epigenetics Group author list**

Eilis Hannon1

Diana Schendel2, 6, 15

Christine Ladd-Acosta3,4

Jakob Grove5,6,7,8,

Christine Søholm Hansen6,9,10

David Michael Hougaard6,9

Michaeline Bresnahan11

Ole Mors6,13

Mads Vilhelm Hollegaard6,9ˆ

Marie Bækvad-Hansen6,9

Mady Hornig11,12

Preben Bo Mortensen6,14,15,16,

Anders D. Børglum5,6,7

Thomas Werge6,10,17,

Marianne Giørtz Pedersen6,13,16,

Merete Nordentoft6,18

Joseph Buxbaum19

M. Daniele Fallin4, 21

Jonas Bybjerg-Grauholm6,9,

Abraham Reichenberg19

Jonathan Mill1

Anna Starnawska 5,6,7,13

Nicklas Heine Staunstrup 5,6,7,13

Magdalena Janecka affiliations 19

Henriette Thisted Horsdal 6, 15

Shantel Weinsheimer6, 10

1 University of Exeter Medical School, University of Exeter, RILD Building, Level 4, Barrack Rd, Exeter EX2 5DW, UK.

2 Department of Public Health, Aarhus University, Aarhus, Denmark.

3 Department of Epidemiology, Johns Hopkins Bloomberg School of Public Health, Baltimore, MD, USA.

4 Wendy Klag Center for Autism and Developmental Disabilities, Johns Hopkins Bloomberg School of Public Health, Baltimore, MD, USA.

5 Department of Biomedicine and Centre for Integrative Sequencing, iSEQ, Aarhus University, Aarhus, Denmark.

6 iPSYCH, The Lundbeck Foundation Initiative for Integrative Psychiatric Research, Aarhus, Denmark.

7 Centre for Genomics and Personalized Medicine, Aarhus, Denmark.

8 Bioinformatics Research Centre, Aarhus University, Aarhus, Denmark.

9 Center for Neonatal Screening, Department for Congenital Disorders, Statens Serum Institut, Copenhagen, Denmark.

10Institute of Biological Psychiatry, MHC Sct. Hans, Mental Health Services Copenhagen, Roskilde, Denmark.

11Center for Infection and Immunity, Columbia University Mailman School of Public Health, New York, USA.

12Department of Epidemiology, Columbia University Mailman School of Public Health, New York, USA.

13Psychosis Research Unit, Aarhus University Hospital, Risskov, Denmark.

14Department of Clinical Medicine, Aarhus University; Aarhus University Hospital, Risskov, Denmark.

15National Centre for Register-Based Research, Aarhus University, Aarhus, Denmark.

16Centre for Integrated Register-based Research, Aarhus University, Aarhus, Denmark.

17Department of Clinical Medicine, University of Copenhagen, Copenhagen, Denmark.

18Mental Health Services in the Capital Region of Denmark, Mental Health Center Copenhagen, University of Copenhagen, Copenhagen, Denmark.

19 Icahn School of Medicine at Mount Sinai, New York, NY, USA

20Department of Psychiatry, Columbia University, New York, USA.

21Department of Mental Health, Johns Hopkins Bloomberg School of Public Health, Baltimore, MD, USA.

**Supplementary Figure 1: Manhattan plot and qq-plot of the MWAS for the CCC scores**

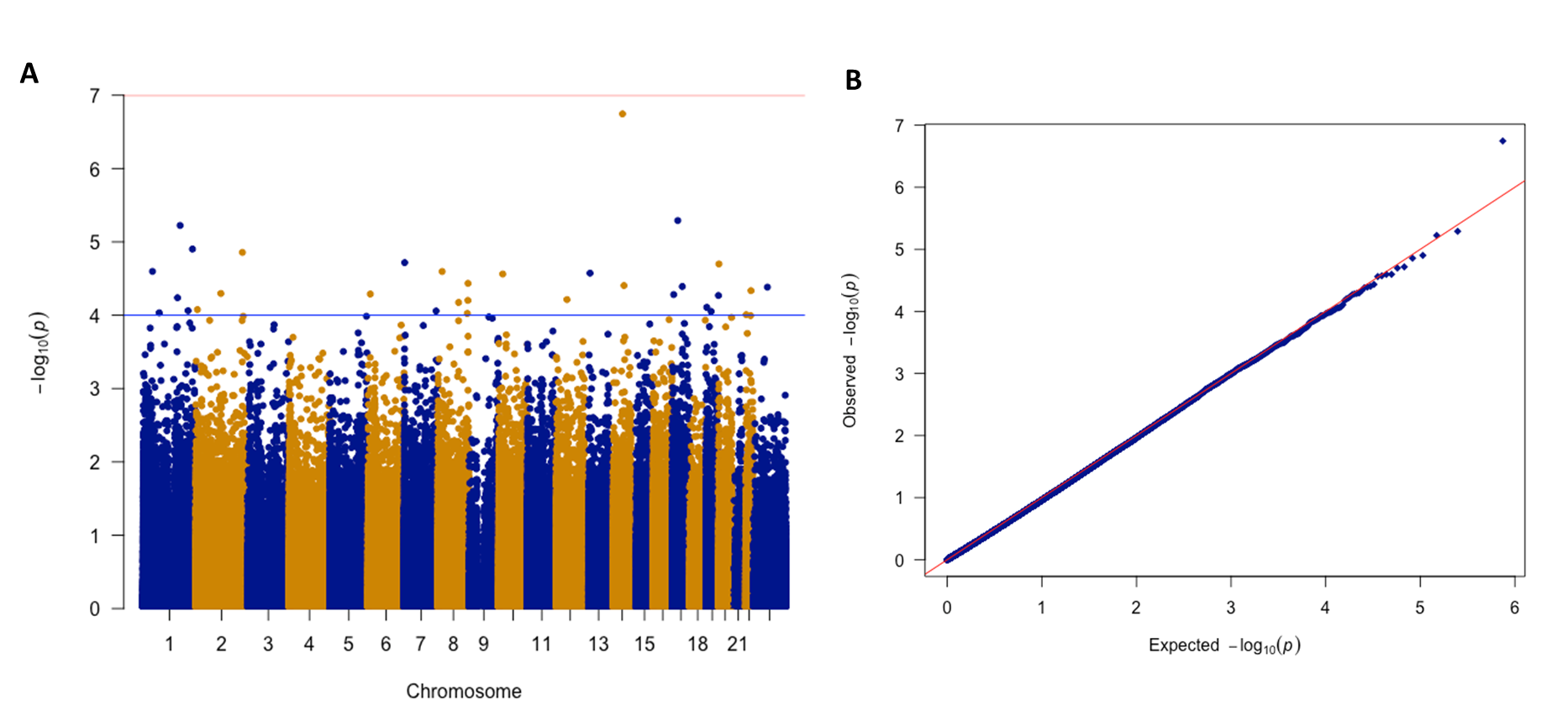

*A: Manhattan plot of the CCC MWAS. B: QQ plot of the CCC MWAS*

**Supplementary Table 1: List of CpGs with P < 0.001 in the SCDC MWAS**

| Name | Chromosome | Position | Gene symbol | Gene Region | BETA | SE | P |
| --- | --- | --- | --- | --- | --- | --- | --- |
| cg00156802 | chrX | 19358457 | chrX:19361726-19362479 |  | -0.81768 | 0.2024 | 5.35E-05 |
| cg03098447 | chr17 | 7210075 | EIF5A;EIF5A;EIF5A | TSS1500;TSS1500;TSS1500 | 2.499287 | 0.641473 | 9.77E-05 |
| cg03202738 | chr4 | 1.56E+08 | NPY2R | TSS200 | 3.25216 | 0.791181 | 3.95E-05 |
| cg05125693 | chr10 | 94051830 | CPEB3;MARCH5 | TSS1500;Body | 2.865308 | 0.724334 | 7.63E-05 |
| cg05877109 | chr14 | 23775794 | BCL2L2 | TSS1500 | 2.574715 | 0.583753 | 1.03E-05 |
| cg07640800 | chr2 | 80529315 | LRRTM1;CTNNA2;CTNNA2 | 3'UTR;Body;Body | 1.673823 | 0.381845 | 1.17E-05 |
| cg10894566 | chr15 | 89905901 | chr15:89904822-89906050 | | 5.931252 | 1.379222 | 1.70E-05 |
| cg11228785 | chr5 | 1.79E+08 | ADAMTS2;ADAMTS2 | Body;Body | 1.080653 | 0.275089 | 8.55E-05 |
| cg11416605 | chrX | 63267928 | chrX:63263904-63264129 |  | 1.233394 | 0.296407 | 3.17E-05 |
| cg11490681 | chr17 | 77460916 | HRNBP3 | 5'UTR | 0.591558 | 0.151996 | 9.94E-05 |
| cg13448605 | chr1 | 38100467 | RSPO1;RSPO1 | 1stExon;5'UTR | 11.17919 | 2.843069 | 8.42E-05 |
| cg14379490 | chr9 | 96221055 | FAM120A | Body | -1.78917 | 0.35686 | 5.34E-07 |
| cg15925695 | chr11 | 1.12E+08 | DIXDC1;DIXDC1 | TSS200;Body | 2.998837 | 0.745251 | 5.72E-05 |
| cg17185953 | chr1 | 53387522 | ECHDC2 | TSS200 | 3.780007 | 0.964301 | 8.86E-05 |
| cg19478343 | chr20 | 49620679 | KCNG1 | Body | -1.6772 | 0.402575 | 3.10E-05 |
| cg19984781 | chr6 | 30710898 | FLOT1 | TSS1500 | 0.932476 | 0.237908 | 8.87E-05 |
| cg25165908 | chr12 | 1.11E+08 | CUX2 | Body | -1.17914 | 0.299162 | 8.10E-05 |
| cg25377985 | chr4 | 1.48E+08 | TTC29 | TSS1500 | 1.876305 | 0.464107 | 5.28E-05 |
| cg27314761 | chr2 | 1.97E+08 | SLC39A10;SLC39A10 | 5'UTR;5'UTR | -1.07153 | 0.248946 | 1.68E-05 |
